# Sensory representations and pupil-indexed listening effort provide complementary contributions to multi-talker speech intelligibility

**DOI:** 10.1101/2023.08.13.553131

**Authors:** Jacie R. McHaney, Kenneth E. Hancock, Daniel B. Polley, Aravindakshan Parthasarathy

## Abstract

Optimal speech perception in noise requires successful separation of the target speech stream from multiple competing background speech streams. The ability to segregate these competing speech streams depends on the fidelity of bottom-up neural representations of sensory information in the auditory system and top-down influences of effortful listening. Here, we use objective neurophysiological measures of bottom-up temporal processing using envelope-following responses (EFRs) to amplitude modulated tones and investigate their interactions with pupil-indexed listening effort, as it relates to performance on the Quick speech in noise (QuickSIN) test in young adult listeners with clinically normal hearing thresholds. We developed an approach using ear-canal electrodes and adjusting electrode montages for modulation rate ranges, which extended the rage of reliable EFR measurements as high as 1024Hz. Pupillary responses revealed changes in listening effort at the two most difficult signal-to-noise ratios (SNR), but behavioral deficits at the hardest SNR only. Neither pupil-indexed listening effort nor the slope of the EFR decay function independently related to QuickSIN performance. However, a linear model using the combination of EFRs and pupil metrics significantly explained variance in QuickSIN performance. These results suggest a synergistic interaction between bottom-up sensory coding and top-down measures of listening effort as it relates to speech perception in noise. These findings can inform the development of next-generation tests for hearing deficits in listeners with normal-hearing thresholds that incorporates a multi-dimensional approach to understanding speech intelligibility deficits.

## Introduction

Everyday listening in multi-talker environments involves a complex interplay between the neural encoding of acoustic information, and the top-down influences of cognitive factors that form a part of effortful listening. Current clinical assessments of hearing impairments are inadequate in assessing this inherently multi-dimensional process because they place heavy emphasis on peripheral auditory function and far less on the sequelae of neural coding that follows cochlear transduction. The standard audiological battery is not sensitive enough to capture hearing difficulties reported by 5-10% of patients seeking help at the audiology clinic, and yet present with normal audiograms^1,2^.

Deficiencies in neural encoding of afferent sensory information within the auditory pathway reflects a complex mixture of degeneration and compensation at successive stations of auditory processing. Sensory degenerations, such as the loss of outer hair cell or strial function can be measured by an audiogram. However, cochlear neural degenerations caused by the loss of synapses between the inner hair cells and the auditory nerve are ubiquitous but remain ‘hidden’ to current clinical tests^3–7^. Peripheral deafferentation is often accompanied by compensatory plasticity, or a relative increase of activity in central auditory structures^8–11^. Although increased central ‘gain’ may benefit listening in quiet, it is likely maladaptive for listening in noise^12^.

In animal models, changes in peripheral neural encoding and resultant compensatory gain can be measured by directly assessing relative neural activity in ascending auditory structures^8,11–13^. However, in humans, indirect measurements of central gain is typically achieved using auditory evoked potentials - ensemble neural activity to sound recorded at the scalp^14^. One such auditory evoked potential is the envelope following response (EFR)—steady-state potentials evoked by the neural synchronization, or phase-locking, to the stimulus amplitude envelope. The phase-locking abilities of neurons are biophysically constrained and decrease along the ascending auditory pathway. Specifically, the upper limit of phase-locking at cortical regions of the auditory pathway are ∼80Hz, while phase-locking limits of the auditory nerve exceed ∼1000Hz (see for review^15^). By exploiting these divergent phase-locking limits of the auditory pathway, EFRs can emphasize cortical, midbrain, and brainstem sources by altering the temporal modulation of the stimulus amplitude that is used to drive the response^6,16–20^. Hence, by comparing EFRs to fast rates and EFRs to slower rates, one can ideally obtain a picture of auditory temporal processing abilities along the entire auditory neuraxis.

In addition to the fidelity of sensory encoding, multi-talker speech intelligibility involves the recruitment of additional cognitive resources that are diverted to assist with listening^21^. This intentional reallocation of cognitive resources, broadly referred to as listening effort, can be indexed using task-related changes in pupil diameter^21–23^. Increase in task-linked pupil diameter has been linked with general cognitive functioning, task difficulty, arousal, and speech intelligibility^24–29^. We have previously measured pupil-indexed effort while subjects identified spoken streams of monosyllabic digits devoid of linguistic context, in the presence of multiple competing digit streams^2^. Changes in pupil diameter were modulated by task difficulty and related to behavioral outcomes. However, the extent to which effort changes with added linguistic context, and the extent to which these top-down processes interact with sensory encoding in the auditory pathway remains unclear.

In the current study, we demonstrate that EFRs can be reliably measured to a variety of modulation frequencies to emphasize complementary neural generators along the ascending auditory pathway in a cohort of young adults with normal hearing thresholds. Further, pupil-indexed listening effort was modulated by task difficulty in subjects performing the Quick Speech in Noise test (QuickSIN^30^), a clinically relevant multitasker speech intelligibility task with moderate linguistic and contextual cues. Finally, using a multivariate regression model, we show that bottom-up sensory coding and top-down listening effort provide complementary contributions to multi-talker intelligibility.

## Methods

### Participants

Nineteen English-speaking participants (mean age = 28.70, *SD* = 4.15 years) were recruited from the greater Boston, MA area to complete all portions of the study. Three participants did not complete the electrophysiological portion of the experiment, resulting in a final sample size of sixteen (6 male, 10 female). Participant eligibility was determined on the first visit by screening for cognitive skills (Montreal Cognitive Assessment > 25^31^), depression (Beck’s Depression Inventory < 21^32^), and tinnitus (Tinnitus reaction questionnaire < 72^33^). All participants had normal hearing sensitivity with thresholds ≤ 25dB for octave frequencies between 250-8000 Hz and were not users of assistive listening devices. Participants received monetary compensation per hour for their participation. This research protocol was approved by the Institutional Review Board at Massachusetts Eye and Ear Infirmary (Protocol #1006581) and Partners Healthcare (Protocol #2019P002423).

### Pupillometry

#### Stimuli and acquisition

Pupillary responses were recorded with a head mounted pupillometry system at a 30 Hz sampling rate (Argus Science ET-Mobile) while participants completed the Quick Speech in Noise (QuickSIN) test^30^. QuickSIN is a standard test of speech perception in noise that provides a measure of signal-to-noise ratio (SNR) loss, which indicates the lowest SNR level at which the listener can accurately identify words 50% of the time. The tests were administered using a Windows Surface tablet. Each QuickSIN test list consisted of six sentences masked in four-talker babble at the following SNR levels: 25, 20, 15, 10, 5, and 0 dB. Sentences were presented in descending order based on SNR level. All participants completed two practice QuickSIN lists before completing four test lists. Participants were instructed to fixate on a point on the screen during listening and to repeat the target sentence to the best of their ability. Each target sentence contained five keywords for identification. The number of key words identified per sentence were recorded. Then, the proportion of keywords correctly identified for each SNR across all four test lists (20 total key words per SNR) was calculated for each participant. Prior to QuickSIN testing, the dynamic range of the pupil was first characterized in each participant by presenting alternating dark and bright screens. The ambient lighting and screen brightness intensity were then adjusted to obtain a baseline pupil size in the middle of the dynamic range.

#### Pupillometry processing and analysis

Pupillary responses time-locked to the onset of the QuickSIN sentences were processed to account for blinks. Blinks were linearly interpolated from approximately 120 ms before to 120 ms after the detected blinks^34,35^. Any trial containing blinks that were longer than 600 ms were removed from further analysis. The pupillary responses were then averaged across all four test lists at each SNR level for each participant. We then excluded SNR 25 from all further analyses, as visual inspection of the averaged pupillary responses at SNR 25 showed different temporal dynamics compared to pupillary responses for the other SNRs. The difference in temporal dynamics at SNR 25 was likely due to the time related to familiarization for the task in each block, as SNR 25 was always presented first in each list. Pupillary responses from 930 ms to 7530 ms time-locked to the onset of the four-talker babble were analyzed for the remaining SNR levels. This time window was selected to encapsulate changes in pupillary responses starting from when the responses diverged from baseline (∼930 ms) and ending at stimulus offset, before the participants moved to respond.

A growth curve analysis (GCA; (Mirman, 2014) was used to obtain a measure of the slope of the pupillary response during listening. GCA uses orthogonal polynomial time terms to model distinct functional forms of the pupillary response over time. A GCA was fit using a first-order orthogonal polynomial to model the interaction with SNR level. This first-order model provided two parameters to explain the pupillary response. The first is the intercept, which refers to the overall change in the pupillary response over the time-window of interest. The second is the linear term (ot1), which represents the slope of the pupillary response over time. The GCA model included fixed effects of SNR (20, 15, 10, 5, and 0; reference = 0) on the linear term with a random slope of the interaction between participant and SNR on the linear time term:

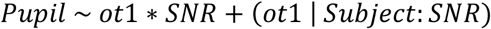

GCA were conducted in R^36^ using the *lme4* package^37^, and *p*-values were estimated using the *lmerTest* package^38^. Multiple comparisons were performed using the *emmeans* package. Adjusted *p*-values are reported using Benjamini-Hochberg Procedure to control for the false discovery rate^39^.

### Electrophysiology

#### Stimuli and acquisition

Electroencephalography (EEG) for the EFRs was collected in an electrically shielded sound attenuating chamber. Participants were seated in a reclined chair and were instructed to minimize movements. The recording session lasted approximately three hours and participants were given breaks as necessary. Recordings were collected using a 16-channel EEG system (Processor: RZ6, preamplifier: RA16PA, differential low impedance amplifier: RA16-LID, TDT System) with two gold-foil tiptrodes positioned in the ear canals (Etymotic) with a sampling rate of 24,414.0625 Hz. Cup electrodes were placed at the Fz site and at both ear lobes, all referenced to a ground at the nape. Electrode impedances were below 1 kΩ after prepping the skin (NuPrep, Weaver and Co.) and applying a conductive gel between the electrode and the skin (EC2, Natus medical).

Envelope following responses (EFRs) were recorded to amplitude modulated (AM) tones. The stimulus carrier frequency was 3000 Hz with amplitude modulation rates of 40, 110, 512, and 1024 Hz. The 3000Hz carrier frequency was chosen to allow the use of modulation frequencies up to 1024 Hz, while remaining within frequencies considered to be relevant to everyday communication. Stimuli were 200 ms long, presented with a 3.1 per second repetition rate in alternating positive and negative polarities. A calibrated ER3A insert earphone in the right ear was used for stimulus presentation. Stimuli presentation level was 85 dB SPL. Stimulus delivery (sampling rate: ∼100 kHz) and signal acquisition were coordinated using the TDT system low impedance amplifier and a presentation and acquisition software (LabVIEW).

#### Electrophysiology processing and analysis

EFRs were processed using a fourth-order Butterworth filter with a lowpass filter of 3000 Hz. The following highpass filter cutoffs were used for 40 Hz, 110 Hz, 512 Hz, and 1024 Hz AM stimuli, respectively: 5 Hz, 80 Hz, 200 Hz, 300 Hz. Fast Fourier transforms (FFTs) were performed on the averaged time domain waveforms for each participant at each AM rate starting 10 ms after stimulus onset to exclude ABRs and ending 10ms after stimulus offset using MATLAB v2022a (MathWorks Inc., Natick, Massachusetts). The maximum amplitude of the FFT peak at one of three adjacent bins (∼3Hz) around the modulation frequency of the AM rate is reported as the EFR amplitude. FFT amplitudes at ten frequency bins on either side of the peak 3 bins were averaged to calculate the noise floor. The signal to noise ratio was calculated as the ratio of the max FFT amplitude at one of the three center bins to the noise floor, expressed in dB.

A GCA was then fit to model EFR amplitudes across each AM rate. The best-fit model that promoted model convergence and did not produce a singular fit contained a first-order orthogonal polynomial with a random slope of the linear time term (ot1) per participant that removed the correlation between the random effects:

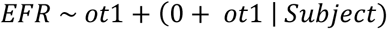

### Statistical Analysis

Correlations between QuickSIN scores with EFR metrics and pupillary metrics were assessed using Spearman’s rank correlations. Spearman’s rank correlations were chosen because Pearson’s correlations are not recommended for sample sizes less than 25^40^. To examine speech intelligibility scores, a linear mixed effects model was fit with QuickSIN score as the outcome variable, a fixed effect of SNR level, and a random effect of participant. Multiple comparisons between SNR levels were performed using the *emmeans* package in R^36^. Adjusted *p*-values are reported using the Benjamini-Hochberg Procedure to control for the false discovery rate^39^.

A backward stepwise multivariate regression was performed to explain variability in QuickSIN at SNR 0 due to EFRs and pupillary responses. The outcome variable was QuickSIN performance at SNR 0, with the pupillary slope at SNR 0 and the EFR slope as predictor variables. The stepwise regression selected the best-fit model based on AIC and adjusted-R^2^ for each model that was calculated. The linear regression was performed in R using *lme4* package^37^. The stepwise regression was performed using the *MASS* package ^41^.

## Results

### EFRs Can be Reliably Measured to Assess Auditory Temporal Processing for Modulation Frequencies up to 1024 Hz in Humans

We first examined the extent to which EFRs could be reliably recorded for the range of modulation frequencies used in this study. EFR amplitudes exhibit a low-pass shape as modulation frequency increases, such that the temporal modulation transfer function decreases logarithmically. Yet, studies in animal models suggest that EFRs to modulation frequencies in the 500-1000Hz AM region can still be recorded reliably above the noise floor^6,20^. Here, we sought to determine if the same was true for humans with our recording setup. **Figure 1** shows the AM stimuli (Fig. 1A), the grand averaged EFR responses in the time domain (Fig. 1B) and the grand averaged FFT spectra of these EFRs in the frequency domain (Fig. 1C). All four modulation frequencies used in this study exhibited robust EFRs, observed both in the time domain, and more clearly, in the frequency domain. In the frequency domain the peaks at modulation frequency were significantly above the noise floor. A quantification of the signal to noise ratio (SNR) revealed an average SNR of approximately 6 dB *(M* = 6.252 dB, *SD* = 0.850 dB) across all modulation frequencies tested (Fig. 1D), suggesting that we can reliably record EFRs within this range in our participant population.

**Figure 1.**
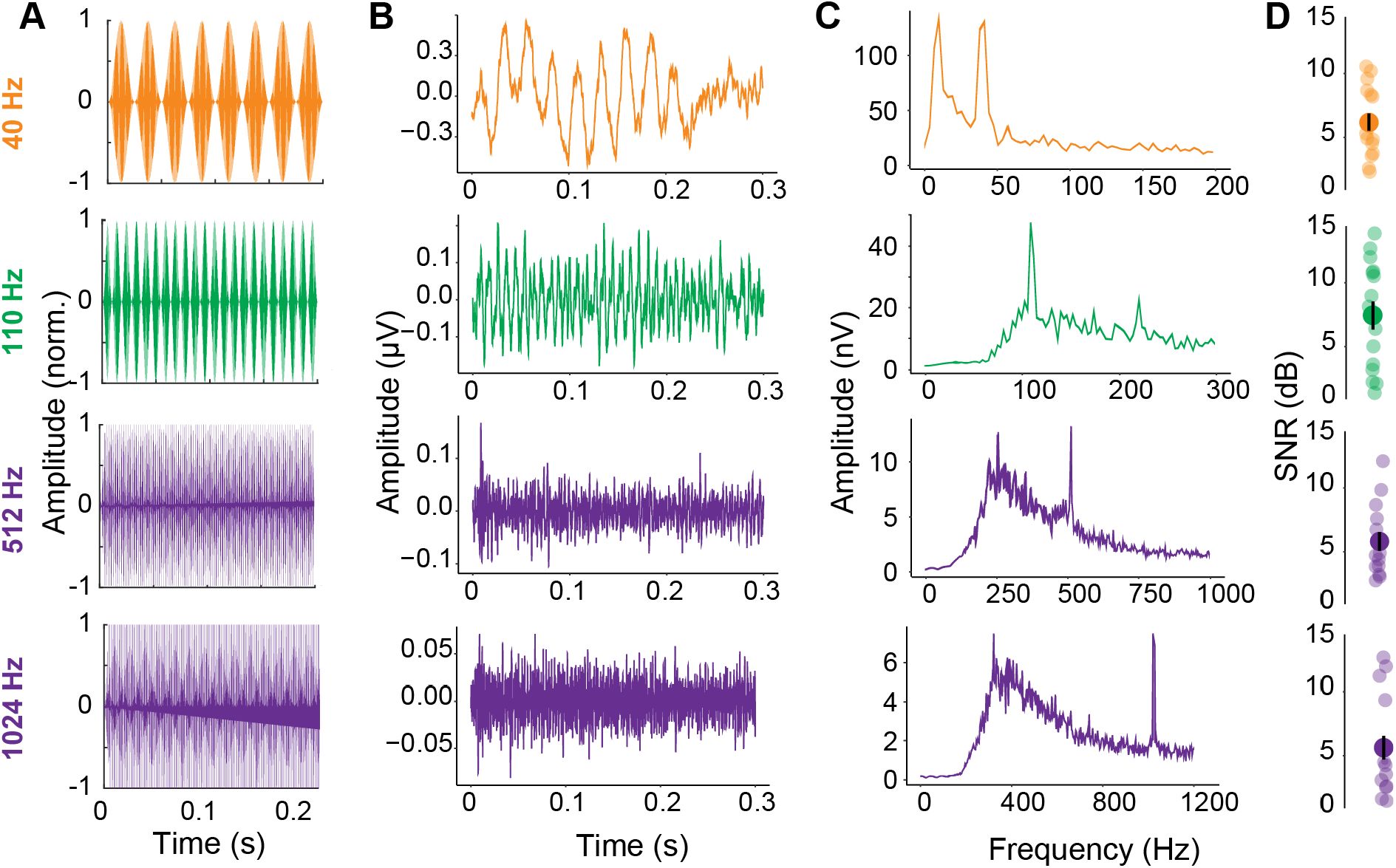
EFRs can be reliably recorded up to modulation frequencies of 1024Hz. **(A)** Time domain stimulus waveforms of the AM tones used in this study. The carrier used was 3 kHz, the modulation depth was 100% and the AM rates were 40Hz, 110Hz, 512Hz and 1024Hz. **(B)** Time domain grand averaged EFRs for each modulation frequency collected for all participants from this study. The neural response ‘follows’ the stimulus amplitude envelope shown in A, most clearly seen for the slower modulation frequencies. **(C)** Responses in (B) transformed to the frequency domain using a fast-Fourier transform exhibits clear peaks at all modulation frequencies tested in this study. **(D)** Signal to noise ratio of EFR peaks in the frequency domain for all participants and all modulation frequencies suggests an average SNR of 6 dB across all modulation frequencies.

### Multi-channel acquisition suggests that EFRs can be utilized to evaluate the relative neural activity from peripheral and central auditory regions

We then sought to confirm that EFRs to various modulation frequencies do in fact emphasize complementary neural generators along the auditory pathway, by leveraging our multichannel approach and comparing EFRs across various electrode montages. The schematic for the hypothesized rationale is displayed in **Figure 2A**. Neural generators are indicated by circles whose sizes correspond to size of the auditory nuclei, with cortical generators having the largest size (orange), and peripheral generators having the smallest (purple). The three electrode montages we compared EFRs across were: 1) Fz to ipsilateral tiptrode placed in the stimulated ear (Fz-R), 2) ipsilateral tiptrode to the contralateral tiptrode placed in the unstimulated ear (L-R), and 3) Fz to contralateral tiptrode (Fz-L). The Fz-R montage should capture contributions from all auditory generators along the ascending pathway^19,42^. The L-R montage should capture peripheral generators while de-emphasizing cortical generators due to the distance the volume conducted signal needs to pass through to be captured by the electrode^42,43^. The Fz-L montage should capture central generators while de-emphasizing peripheral generators for the same reason above. This hypothesis was supported by the comparative EFRs shown in Figure 2B, plotted both as absolute amplitude (left column) and relative change in amplitude compared to the Fz-R condition (right column). EFRs to 40Hz AM had the highest response amplitudes in the Fz-R montage, relative to Fz-L (β = - .047, *t* = -2.785, *p* = .009) and L-R montages (β = -.081, *t* = -4.769, *p* < .001). Response amplitudes significantly decreased by about 30% in the Fz-L montage (*M* = 0.110, *SD* = 0.061) relative to the Fz-R montage (*M* = 0.157, *SD* = 0.083), presumably due to the loss of contributions from subcortical areas. Response amplitudes were further reduced by approximately 40% in the L-R horizontal montage (*M* = 0.076, *SD* = 0.044) relative to the Fz-R montage. At 110 Hz AM rate, EFR amplitudes at Fz-R did not significantly differ from EFR amplitudes in the L-R montage (β = .009, *t* = 1.329, *p* = .194) nor Fz-L montage (β = .004, *t* = 0.558, *p* = .581). The similar EFR amplitudes at 110 Hz across montages was consistent with the idea that ∼110Hz AM reflects a mixture of contributions from cortical and subcortical generators^44,45^, and as such would not differ significantly between electrode montages. EFRs to faster AM rates (i.e., 512 and 1024 Hz) demonstrated changes in amplitudes consistent with generators originating from more peripheral neural sources. EFR amplitudes for 512 Hz AM rate were significantly higher in the L-R montage (β = .006, *t* = 3.314, *p* = .003) relative to the Fz-R montage, though EFR amplitudes did not differ between Fz-R and Fz-L montages (β = -.003, *t* = -1.318, *p* = .198). Trends seen at 512Hz were amplified for 1024 Hz AM, where EFR amplitudes in the Fz-R montage were greater than Fz-L montage (β = -.004, *t* = -5.139, *p* < .001) but did not differ from amplitudes in the L-R montage (β < .001, *t* = 0.285, *p* = .777), suggesting more distal neural generators in the peripheral auditory pathway.

**Figure 2.**
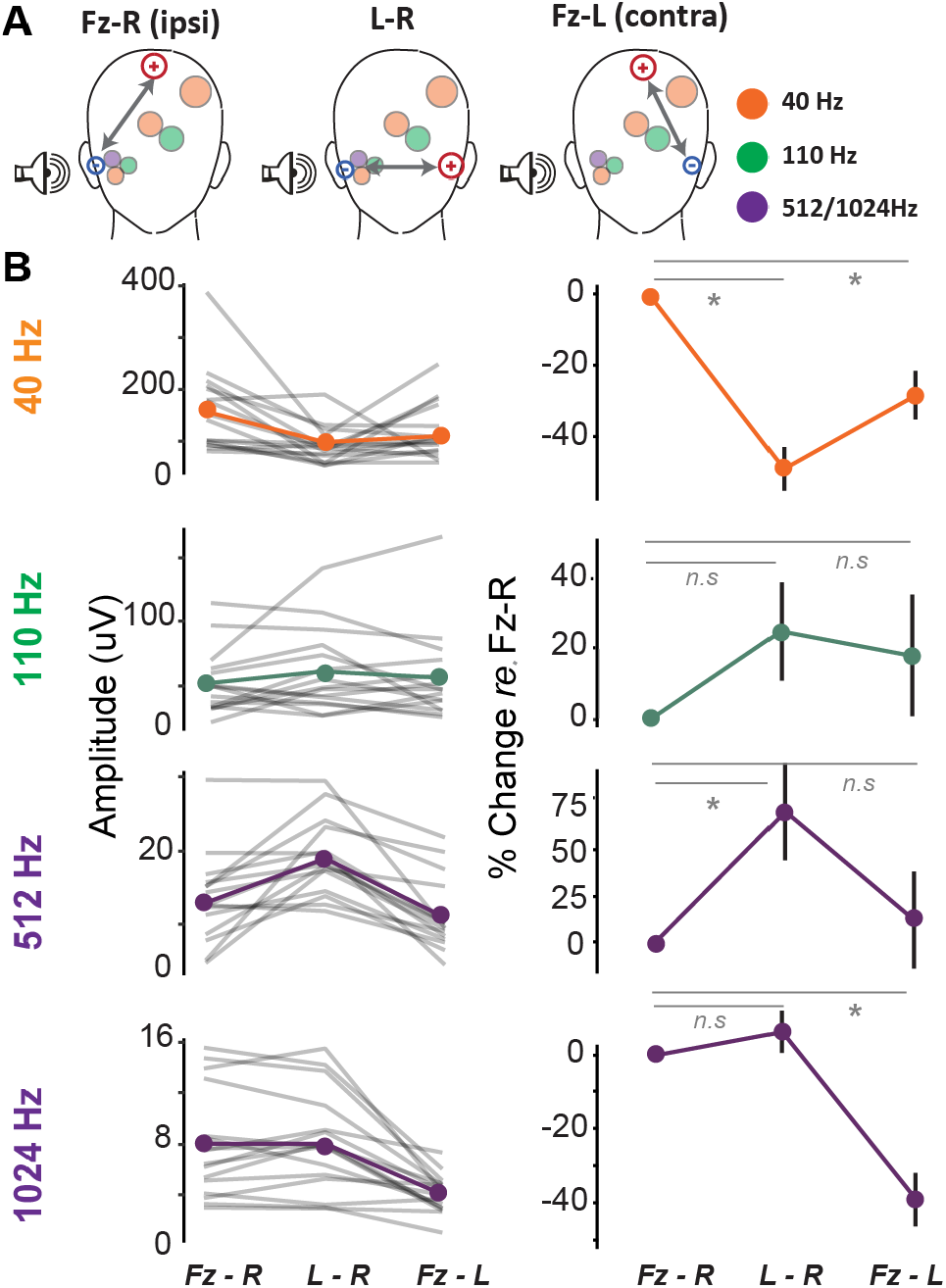
Electrode montage configurations emphasize complementary neural generators. A) Schematic of active and reference electrode placement of three montage configurations: Fz to Right-tiptrode (Fz-R), Left-tiptrode to Right-tiptrode (L-R), and Fz to Left-tiptrode (Fz-L). The shaded circles represent the auditory nuclei that generate these responses, with circle size reflecting the size of the generators. B) Left: Average envelope following responses (EFR) amplitudes for each AM rate and each montage configuration. Right: The percent change in EFR amplitude at each AM rate for L-R and Fz-L configurations, relative to Fz-R.

Taken together, these results suggest that EFRs to varying AM rates emphasize peripheral versus central neural generators. Further, the Fz-R montage was best suited to capture EFRs along the entire auditory neuraxis for all AM rates.

### Young listeners with normal audiometric thresholds exhibited substantial variability in both EFR amplitudes and speech perception in noise

Participants demonstrated individual differences in EFR amplitudes in the Fz-R montage and the shape of the temporal modulation transfer function (i.e., EFR amplitudes as a function of AM rate; **Figure 3**A, Table 1). To obtain a single metric of the temporal modulation transfer function per participant, we used a GCA to calculate an EFR slope (Fig. 3B). We then used this EFR slope to compare the EFRs to other measures across this study. Mean centered EFR slopes ranged from -0.142 to 0.389. These data suggested that substantial variability in EFR amplitudes across AM rates remained even in young adults with normal audiometric thresholds. Presumably, this variability in EFR amplitudes reflects individual differences in the encoding of temporal information along the ascending auditory pathway.

**Figure 3.**
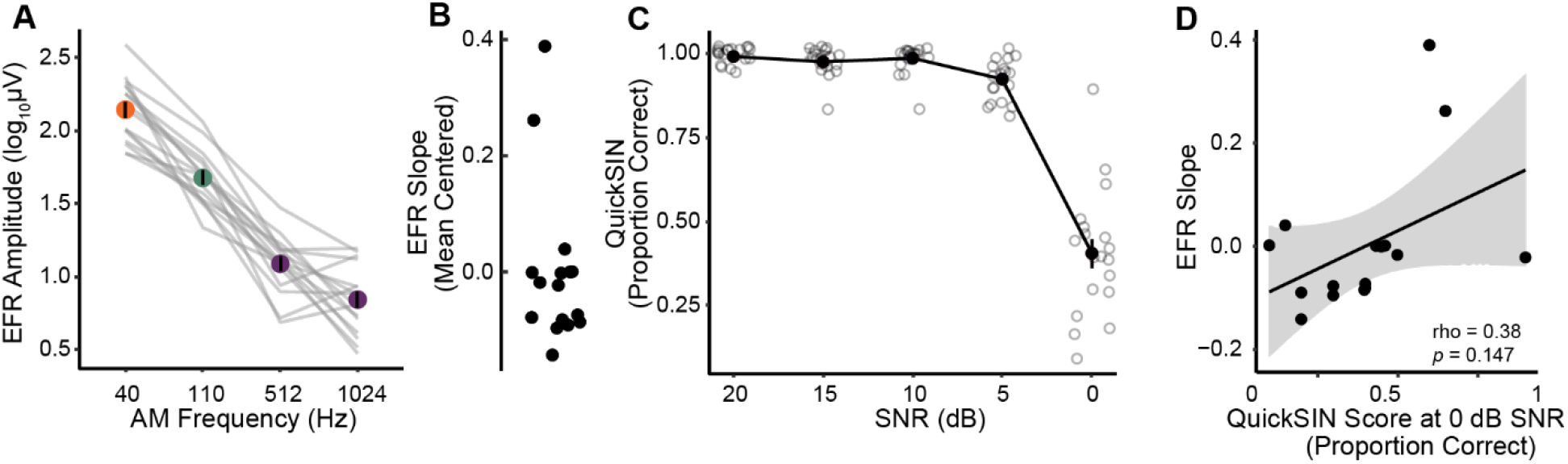
Young listeners with normal audiograms exhibit substantial individual differences in EFRs and speech perception in noise. A) Temporal modulation transfer function with EFR amplitudes at each modulation rate. Individual participant lines are denoted in gray. B) Distribution of individual EFR slope estimates for each participant. C) Proportion of keywords identified correctly across four QuickSIN lists for each SNR level. Solid line and points denote average performance across all participants. Smaller points denote individual participant scores. QuickSIN performance significantly drops from SNR 5 to SNR 0 dB. D) The scatterplot reveals a non-significant correlation between EFR slope estimates and QuickSIN scores at signal-to-noise ratio (SNR) 0 dB.

**Table 1.**
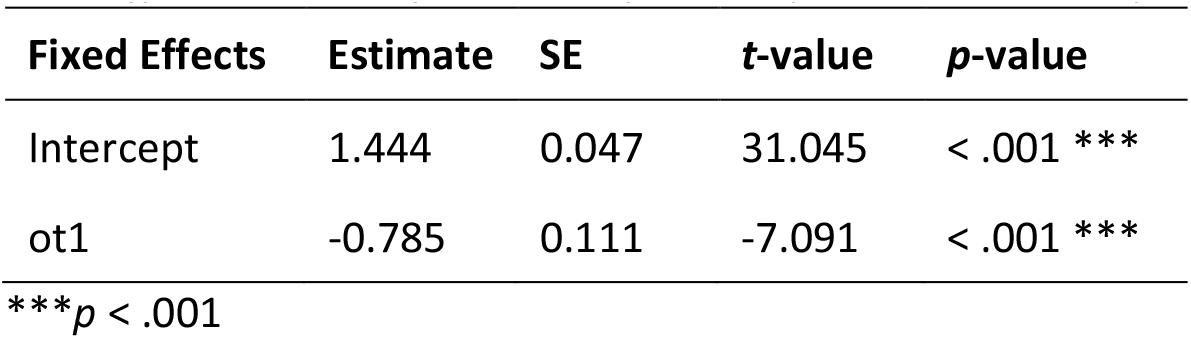
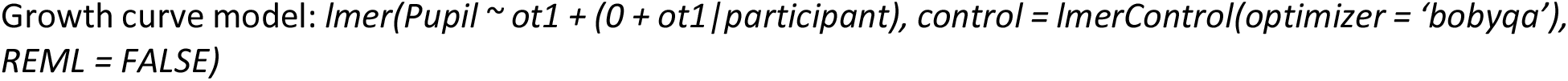
Fixed effect estimates for model of EFR amplitudes across amplitude modulation rates (observations = 62)

We then examined speech perception in noise using QuickSIN, a clinically used multi-talker speech intelligibility task with moderate contextual and linguistic cues (Fig. 3C). Each participant’s speech perception in noise score was calculated as the proportion of correct keywords identified per SNR level across all four test lists. Behavioral performance at SNR 0 dB was significantly lower than performance at the easier SNRs of 20, 15, 10, and 5 dB (*p*s < .001, Linear mixed model, Fig. 3C, Table 2). On average, participants had near-perfect performance for SNRs at 20, 15, 10 and 5dB, ranging from an average of 99% at SNR 20 to an average of 93% at SNR 5. Performance at these SNRs between 20 and 5 dB did not significantly differ from one another (*p*s > .05, Table 2). Performance dropped significantly for most participants at the most challenging SNR of 0 dB, and this drop in performance was statistically significant (ps <0.001, Table 2). Performance at the most challenging SNR also varied widely, with percent correct ranging from 10% to 90% in our participants (*M* = 40.5%, *SD* = 18.8%). These results suggest that while young listeners performed near ceiling at easier listening conditions, there was significant individual variability under challenging listening conditions, despite normal audiometric thresholds.

**Table 2.**
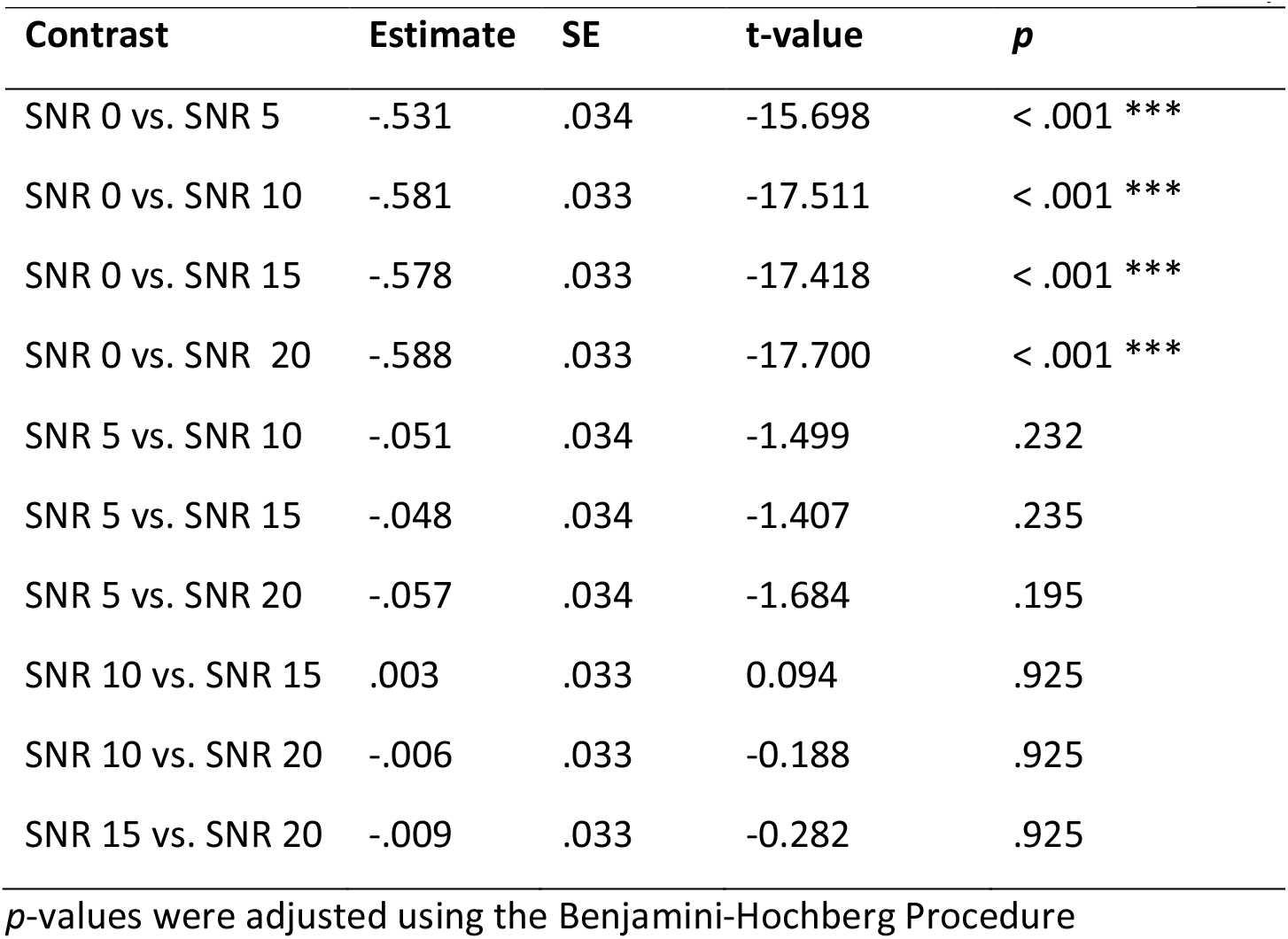
Multiple comparisons from a linear mixed effects model examining QuickSIN performance.

| Contrast | Estimate | SE | t-value | p |
| --- | --- | --- | --- | --- |
| SNR 0 vs. SNR 5 | -.531 | .034 | -15.698 | < .001 *** |
| SNR 0 vs. SNR 10 | -.581 | .033 | -17.511 | < .001 *** |
| SNR 0 vs. SNR 15 | -.578 | .033 | -17.418 | < .001 *** |
| SNR 0 vs. SNR 20 | -.588 | .033 | -17.700 | < .001 *** |
| SNR 5 vs. SNR 10 | -.051 | .034 | -1.499 | .232 |
| SNR 5 vs. SNR 15 | -.048 | .034 | -1.407 | .235 |
| SNR 5 vs. SNR 20 | -.057 | .034 | -1.684 | .195 |
| SNR 10 vs. SNR 15 | .003 | .033 | 0.094 | .925 |
| SNR 10 vs. SNR 20 | -.006 | .033 | -0.188 | .925 |
| SNR 15 vs. SNR 20 | -.009 | .033 | -0.282 | .925 |

*p*-values were adjusted using the Benjamini-Hochberg Procedure

Finally, we explored if there were any correlations between performance on QuickSIN and neural encoding metrics obtained from the EFRs. QuickSIN performance at 0 dB SNR did not significantly correlate with EFR amplitudes at 40 Hz (*rho* = .180, *p* = .496), 110 Hz (*rho* = -.067, *p* = .813), 512 Hz (*rho* = .120, *p* = .678), or 1024 Hz (*rho* = .240, *p* = .380). Further, QuickSIN performance at 0 dB SNR did not significantly correlate with the EFR slope metric (Fig. 3D). These results suggest that there was no direct, linear relationship between these bottom-up measures of sensory encoding and QuickSIN performance within our participant population.

### Increases in listening effort are evident prior to changes in behavioral thresholds

Given the lack of a correlation between the EFRs and performance at SNR 0 on QuickSIN, we then asked whether top-down measures of listening effort changed with SNR and if those changes were more indicative of QuickSIN performance. Isoluminous task related changes in pupil diameters were measured as participants completed the task at various SNR levels (**Figure 4A**). Pupillary responses increased after sound onset and during the presentation of the stimuli. We used a GCA to calculate the change in the pupillary response over time during listening, as well as to obtain a metric of pupillary slope. Pupil diameters were modulated by SNR level, with SNR 0 having the fastest change in pupil diameters in our study population (Table 3; Figure 4B). The slope of the pupillary response (ot1) at SNR 0 was significantly steeper than all other SNR levels (*p*s < .05). This suggests that SNR 0 required the greatest amount of listening effort among the SNRs tested.

**Figure 4.**
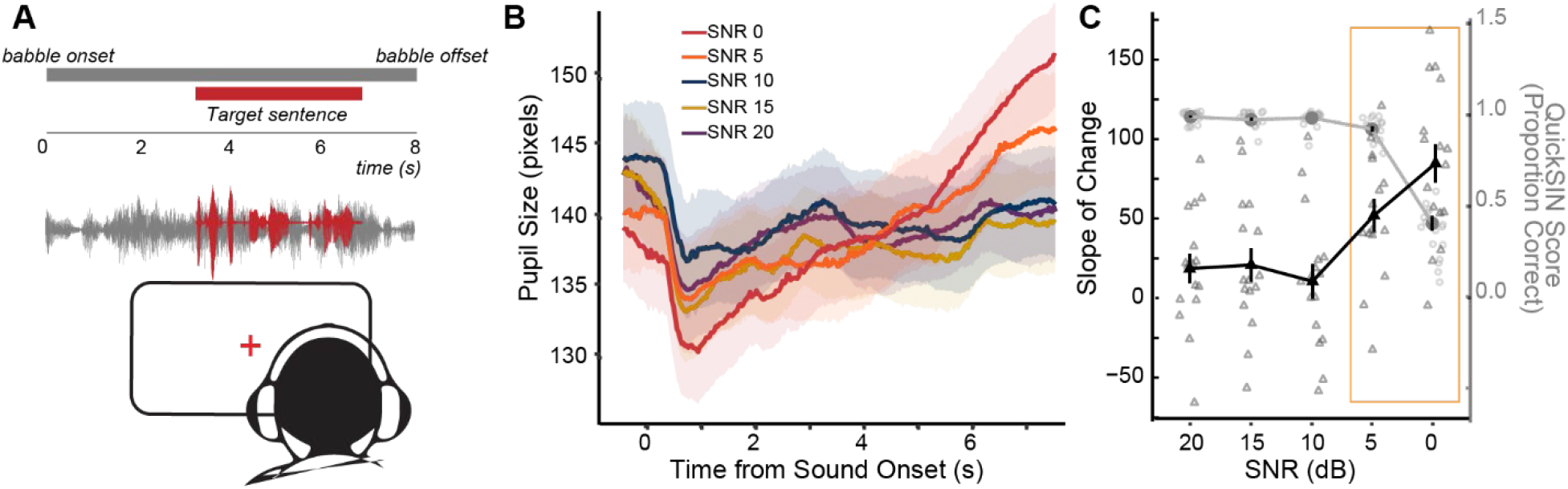
Pupillary responses to QuickSIN show significant changes in listening effort with increasing SNR difficulty. A) Schematic of experimental design. Subjects watched a fixation point and had changes in their pupil diameters measured while listening to QuickSIN sentences. B) Grand-averaged raw pupillary responses to QuickSIN at each SNR level time-locked to the onset of the babble. C) Average QuickSIN performance (left y-axis) at each SNR level in purple, overlaid with the average pupillary slope (right y-axis) at each SNR level in green. Individual participant QuickSIN scores and pupillary slopes are also shown. Listening effort, as measured by the pupillary slopes, at SNR 5 dB significantly increases while QuickSIN performance remains near ceiling, yet the increase in listening effort at SNR 0 dB is not associated with better performance at SNR 0 dB (orange square).

**Table 3.** Multiple comparisons contrasts for the GCA for overall change in the pupillary responses between SNR levels and the slopes of the pupillary responses.

| Contrasts | Estimate | SE | t-value | p-value |
| --- | --- | --- | --- | --- |
| SNR 0 vs. SNR 5 | -0.023 | 9.644 | -0.002 | .998 |
| SNR 0 vs. SNR 10 | 2.147 | 9.487 | 0.226 | .998 |
| SNR 0 vs. SNR 15 | 2.946 | 9.487 | 0.311 | .998 |
| SNR 0 vs. SNR 20 | 2.562 | 9.487 | 0.270 | .998 |
| SNR 5 vs. SNR 10 | 2.170 | 9.644 | 0.225 | .998 |
| SNR 5 vs. SNR 15 | 2.969 | 9.644 | 0.308 | .998 |
| SNR 5 vs. SNR 20 | 2.584 | 9.644 | 0.268 | .998 |
| SNR 10 vs. SNR 15 | 0.799 | 9.487 | 0.084 | .998 |
| SNR 10 vs. SNR 20 | 0.414 | 9.487 | 0.044 | .998 |
| SNR 15 vs. SNR 20 | -0.385 | 9.487 | -0.041 | .998 |
| ot1 x SNR 0 vs. SNR 5 | 0.035 | 0.016 | 2.194 | .047 * |
| ot1 x SNR 0 vs. SNR 10 | 0.078 | 0.015 | 5.055 | < .001 *** |
| ot1 x SNR 0 vs. SNR 15 | 0.067 | 0.015 | 4.356 | < .001 *** |
| ot1 x SNR 0 vs. SNR 20 | 0.070 | 0.015 | 4.509 | < .001 *** |
| ot1 x SNR 5 vs. SNR 10 | 0.044 | 0.016 | 2.779 | .014 * |
| ot1 x SNR 5 vs. SNR 15 | 0.033 | 0.016 | 2.091 | .052 |
| ot1 x SNR 5 vs. SNR 20 | 0.035 | 0.016 | 2.242 | .047 * |
| ot1 x SNR 10 vs. SNR 15 | -0.011 | 0.015 | -0.699 | .606 |
| ot1 x SNR 10 vs. SNR 20 | -0.008 | 0.015 | -0.546 | .650 |
| ot1 x SNR 15 vs. SNR 20 | 0.002 | 0.015 | 0.153 | .878 |

\* $p < .05$ ; \*\*\* $p < .001$
$p$ -values were adjusted using the Benjamini-Hochberg procedure.

Individual participant’s pupillary slopes at each SNR were then extracted from the random effect of the GCA. Overlaying the average pupillary slopes at each SNR against task performance, we found that pupillary slopes were relatively flat for SNRs 20, 15 and 10 dB where task performance was near ceiling (Fig. 4C). Interestingly, pupillary slopes were significantly greater at SNR 5 dB relative to SNR 10 and SNR 20 (Table 3), even though performance was still near ceiling (Table 2). There was an even greater increase in the steepness of the slope at SNR 0, where QuickSIN scores also dropped significantly. While these data suggest that an increase in task difficulty co-occurred with an increase in listening effort as indexed by pupillometry, it is interesting to note that pupillometric changes were not necessarily reflective of behavioral performance outcomes. The increase in the pupillary slope at SNR 5 suggests an increase in listening effort that was enough to maintain near-ceiling performance, but the increase in listening effort at SNR 0 did not counteract the detrimental effects of noise, resulting in a decreased performance relative to easier SNRs.

### Sensory coding measures and listening effort provide complementary contributions to the variance in speech perception in noise

Given the results seen above, we focused further analyses on the most challenging two SNRs – 1) SNR 5 where performance was near-ceiling, but effort increased, and 2) SNR 0 where effort increased but performance dropped. The individual pupillary slopes at SNR 0 were not significantly correlated with task performance at 0 dB SNR (**Figure 5A**) or 5dB SNR (*rho* = -0.310, *p* = 0.212). Additionally, these pupillary responses at SNR 0 were not significantly correlated with EFR amplitudes (40 Hz: *rho* = -.085, *p* = .756; 110 Hz: *rho* = -0.160, *p* = 0.558; 512 Hz: *rho* = -0.190, *p* = 0.490; 1024 Hz: *rho* = -0.490, *p* = 0.054;) or EFR slope (Fig. 5B). These data suggest that there was no direct relationship between the neural encoding measures, listening effort, and multi-talker speech intelligibility within our participant population.

**Figure 5.**
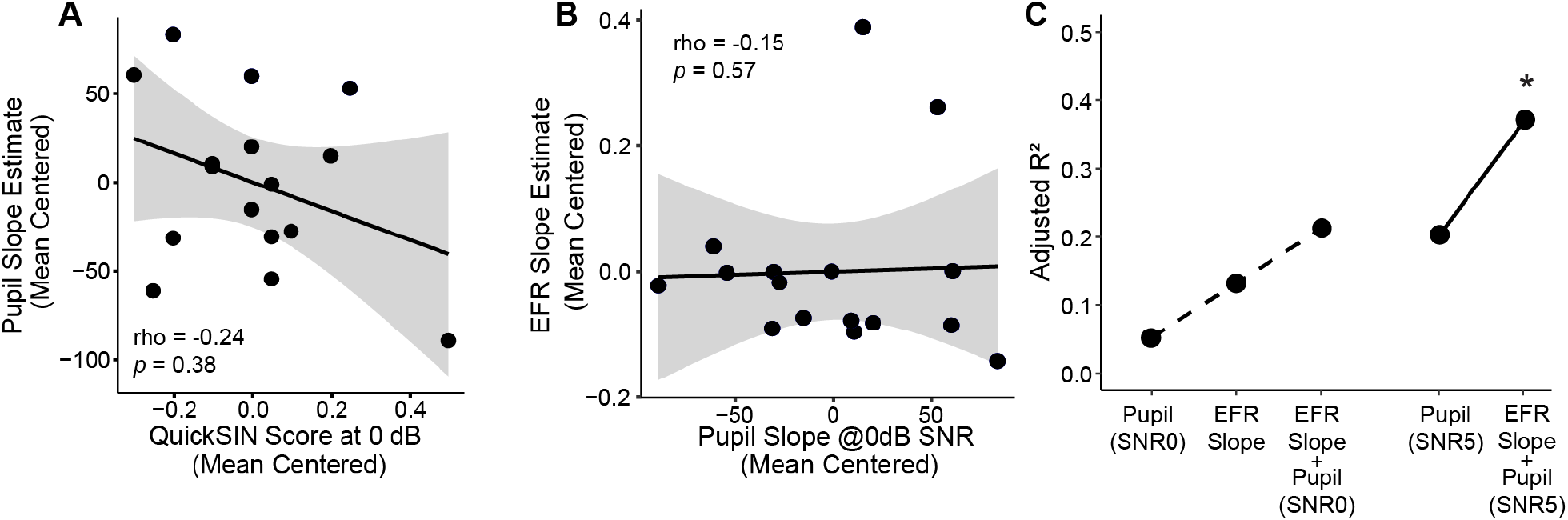
Pupillary slopes and the slope of envelope following responses (EFR) significantly contribute to multi-talker speech intelligibility. **A)** Pupillary slopes at SNR 0 dB (mean centered) are not associated with QuickSIN performance at SNR 0 dB. **B)** Scatterplot between envelope following response (EFR) slopes and pupil slopes at SNR 0 dB. **C)** A stepwise regression measuring the variance explained (Adjusted-R^2^) of QuickSIN at SNR 0 dB revealed incremental improvement with the combination of both EFR slope and pupil slope at SNR 0 dB (dashed line). A second stepwise regression showed an even greater increase in the variance explained in QuickSIN at SNR 0 dB when including the pupil slope at SNR 5 dB (solid line).

To probe the potential synergistic effect between sensory coding and listening effort, we used a backward stepwise regression to calculate the variance explained by EFRs and pupil diameters on performance in the QuickSIN task. For multi-talker speech performance at SNR 0, the stepwise linear regression revealed that a combination of both pupillary slope at SNR 0 and the EFR slope (*SpIN SNR* 0 ∼ *Pupil SNR* 0 + *EFR*) provided the best model fit, with an Adjusted-*R*^2^ = .254 (*F*_(2,13)_ = 3.024, *p* = .084). This suggests that the combination of EFR slope and pupillary slope at SNR 0 explained 21.3% of the variance in QuickSIN performance at 0 dB SNR. Models that included only pupillary slope at SNR 0 or EFR slope explained 5.3% and 13.3% of the variance in QuickSIN at SNR 0, respectively (Fig. 5C).

Interestingly, when the stepwise regression model to explain performance at 0 dB SNR was constructed with just the pupillary slope at SNR 5 instead of SNR 0, the pupillary slope at SNR 5 explained 20.3% of the variance in QuickSIN at SNR 0. The addition of EFR slopes to this model nearly doubled the Adjusted-R^2^ to 37.0%. Additionally, the overall model with EFR slopes and the SNR 5 pupillary slope was significant (*F*_(2,12)_ = 5.109, *p* = .025). Hence, these results suggest that sensory neural encoding and listening effort provided complementary contributions to the overall variance explain in speech perception in noise. Further, the amount of listening effort required to maintain performance at the relatively easier SNR 5 is a better indicator of an individual’s multi-talker speech intelligibility at the more challenging SNR 0.

## Discussion

Speech intelligibility is an inherently multi-dimensional process that includes exquisite interactions between sensory-motor systems, language networks, and top-down attentional networks that include cognition and arousal^46–56^. A combination of influences due to diverse factors such as peripheral hearing thresholds, cochlear health, central auditory processing, arousal state, cognitive status, current emotional status, and familiarity with context of the conversation affects multi-talker speech intelligibility^53,57–63^. Studies that probe the individual effects of these factors by carefully controlling for as many of these factors as possible can only explain a small part of the overall individual variance in speech perception performance. Here, we explore the synergistic contributions of two factors affecting multi-talker speech intelligibility – bottom-up sensory coding fidelity to temporally modulated sounds that was indexed using EFRs and top-down cognitive load that was indexed using pupillometry. We found that although these two factors individually contributed to only a small percentage of the overall variance in our multi-talker speech task, they combined synergistically to explain a significant portion of the variance in performance on our task (Fig. 5C). These results are a first step towards a multi-dimensional approach to understanding speech perception in noise.

EFRs have been used in multiple studies to assess neural coding to the temporally modulated stimulus amplitude envelope^56,64–69^. The neural generators of EFRs evoked to speech sounds, also referred to more generically as the frequency-following response, have been a subject of recent debate. Historically considered to have primarily subcortical generators, recent evidence suggests a larger cortical contribution to these responses^70–74^. In contrast, EFRs evoked to simpler sinusoidally amplitude modulated tones have long been known to emphasize cortical or subcortical generators depending on the AM frequency used, with slower (<40Hz) AM rates thought to emphasize cortical generators, and faster (100-300Hz) AM rates thought to emphasize subcortical generators^16,17,75^. Recent studies also suggest much faster AM rates to be sensitive to peripheral neural degeneration, potentially stemming from the auditory nerve^6,20,76^. These differential neural generators are possibly due to a fundamental biophysical property of the neurons within the auditory pathway, wherein the phase-locking abilities of the auditory nerve neurons extend up to approximately 2000 Hz, but this upper limit gradually decreases along the afferent pathway to about 80Hz in the auditory cortex^15^.

While human studies only use a few modulation rates due to limitations of recording time, animal studies have systematically characterized these EFR generators using lesioning, anesthesia or peripheral nerve damage^6,19,20,75^. Human studies also typically do not use modulation rates faster than ∼300Hz, as the decrease in EFR amplitudes with rate is logarithmic, reaching very small amplitudes at faster rates. Here, through a combination of high sampling rate, ear-canal electrodes, and a low recording noise floor, we demonstrated that we can reliably record EFRs to AM rates up to 1024Hz in young listeners with normal hearing thresholds (Fig. 1). We further compared EFRs across multiple electrode montages to support the idea that faster AM rates emphasize more peripheral generators (Fig. 2). Our findings are in agreement with previous human and animal model studies that used similar multi-channel approaches to determine EFR generators^19,42,43,77–79^. We further used a GCA to determine the slope of EFR change with modulation frequency, providing a metric for assessing temporal processing along the ascending auditory pathway (Fig. 3A-C). This slope metric, and the associated central gain metric also have the added benefit of minimizing inter-subject variability due to recording factors such as head size or electrode impedance, as they are normalized within subject by design.

Individuals showed significant variability in performance on QuickSIN, especially at the most challenging SNR of 0dB, despite being young and having clinically normal hearing thresholds (Fig. 3D). We had previously demonstrated a similar variability in multi-talker speech intelligibility using a digits task consisting of a target speaker and two competing co-localized speakers producing speech streams of a closed set of monosyllabic numbers devoid of linguistic context^2^. Similar results were observed when using QuickSIN, which is a clinically relevant tool for assessing speech perception in noise, consisting of open-set sentences with linguistic context masked in multi-talker babble^30^. Listening effort, assessed using pupillometry, also increased with more challenging SNRs, with SNR 0 resulting in the greatest change in pupil diameter. Pupillometry has rapidly gained prominence as a tool to assess cognitive load and effortful listening, which is modulated by task difficulty, cognitive status and hearing acuity^28,53,80–84^. The precise neural pathways that induce pupillary changes during listening is still under study, but it is hypothesized to be mediated by the locus coeruleus–norepinephrine (LC-NE) system. The LC-NE system is a network of neurons that has wide-spread projections throughout the cortex. Changes in LC-NE system activity are strongly associated with changes in pupil diameter^85^. However, the underlying neural mechanisms modulating pupillary changes may encompass networks that go beyond the LC-NE system, and may be driven by other networks that modulate arousal states^29,86^. Animal studies demonstrate some support for this idea, with pupil diameter indexing momentary changes in arousal states and indicative of task performance under challenging listening conditions^27,85,87^. These studies suggest a non-linear relationship between pupil-indexed arousal and behavior.

Optimal states of arousal result in improved behavioral outcomes. However, pupillary changes that are lower than this optimal state result in disengagement, and changes that are higher than the optimal state result in hyperarousal, both of which affect behavioral performance^27^. This suggests that effortful listening, as indexed by pupillometry, is only valuable to a certain extent. That is, when a listening situation becomes too difficult, increased listening effort is not necessarily beneficial^23^. This non-monotonic relationship can also help to explain our finding wherein listening effort at SNR 5dB explained more variance in behavioral performance at SNR 0 than listening effort at SNR 0 itself. Our finding suggests that an individual who had reached their optimal, intermediate listening effort level at SNR 5 would have likely experienced a decrease in performance at SNR 0. Speech perception at SNR 0 for these individuals was likely deemed to be too difficult, such that an increase in listening effort at SNR 0 did not benefit performance. Conversely, an individual who was approaching the optimal, intermediate listening effort level at SNR 5 would likely experience less of a decrease in performance at SNR 0 compared to someone who already surpassed their optimal performance level by SNR 5.

In our study, neither the EFR slopes, nor the pupillary changes were directly related to QuickSIN task performance. Yet, together, they provided a greater proportion of variance explained compared to either metric in isolation. Pupillary dilations increased significantly for SNR 5, even though there was no added benefit or detriment seen in behavioral performance at that SNR (Fig. 4B). Interestingly, this change in pupil diameter at SNR 5 was a greater indicator for behavioral performance at SNR 0, compared to pupillary changes at SNR 0 itself (Fig. 5C). This suggests that perhaps the ability to expend effort to maintain performance at a less challenging SNR is more predictive of performance at harder SNRs. Future work will explore these interactions between SNRs and pupillary changes, as well as assess if either the sensory coding component or the top-down indices of effortful listening change independently or concurrently with hearing pathologies.

## Author Contributions

AP and DP designed the study. AP collected the data. JM analyzed the data and prepared the initial draft of the manuscript. KEH provided software and support for data acquisition and analysis. All authors contributed to the final version of the manuscript.

## Data availability

Data available upon reasonable request to the corresponding author.

## Acknowledgements

This work was supported by P50DC015817 (DP), T32DC011499 (trainee: JRM), and F31DC020085 (JRM). The authors thank Jessica L’Heureux and Jennifer Klara for administrative and technical support.

## References

1. Hind, S. E. et al. Prevalence of clinical referrals having hearing thresholds within normal limits. International Journal of Audiology 50, 708–716 (2011).

2. Parthasarathy, A., Hancock, K. E., Bennett, K., DeGruttola, V. & Polley, D. B. Bottom-up and top-down neural signatures of disordered multi-talker speech perception in adults with normal hearing. eLife 9, e51419 (2020).

3. Kujawa, S. G. & Liberman, M. C. Adding Insult to Injury: Cochlear Nerve Degeneration after ‘Temporary’ Noise-Induced Hearing Loss. Journal of Neuroscience 29, 14077–14085 (2009).

4. Kujawa, S. G. & Liberman, M. C. Synaptopathy in the noise-exposed and aging cochlea: Primary neural degeneration in acquired sensorineural hearing loss. Hearing Research (ePub ahead of print), (2015).

5. Sergeyenko, Y., Lall, K., Liberman, M. C. & Kujawa, S. G. Age-Related Cochlear Synaptopathy: An Early-Onset Contributor to Auditory Functional Decline. Journal of Neuroscience 33, 13686–13694 (2013).

6. Parthasarathy, A. & Kujawa, S. G. Synaptopathy in the aging cochlea: Characterizing early-neural deficits in auditory temporal envelope processing. The Journal of Neuroscience (2018) doi:10.1523/jneurosci.3240-17.2018.

7. Wu, P. Z. et al. Primary Neural Degeneration in the Human Cochlea: Evidence for Hidden Hearing Loss in the Aging Ear. Neuroscience (2018) doi:10.1016/j.neuroscience.2018.07.053.

8. Chambers, A. R. et al. Central Gain Restores Auditory Processing following Near-Complete Cochlear Denervation. Neuron 89, 867–879 (2016).

9. Auerbach, B. D., Radziwon, K. & Salvi, R. Testing the Central Gain Model: Loudness Growth Correlates with Central Auditory Gain Enhancement in a Rodent Model of Hyperacusis. Neuroscience 407, 93–107 (2019).

10. Parthasarathy, A., Bartlett, E. L. & Kujawa, S. G. Age-related Changes in Neural Coding of Envelope Cues: Peripheral Declines and Central Compensation. Neuroscience 407, 21–31 (2019).

11. Parthasarathy, A., Herrmann, B. & Bartlett, E. L. Aging alters envelope representations of speech-like sounds in the inferior colliculus. Neurobiol. Aging 73, 30–40 (2019).

12. Resnik, J. & Polley, D. B. Cochlear neural degeneration disrupts hearing in background noise by increasing auditory cortex internal noise. Neuron (2021) doi:10.1016/j.neuron.2021.01.015.

13. McGill, M. et al. Neural signatures of auditory hypersensitivity following acoustic trauma. Elife 11, e80015 (2022).

14. Rumschlag, J. A. et al. Age-Related Central Gain with Degraded Neural Synchrony in the Auditory Brainstem of Mice and Humans. Neurobiol Aging 115, 50–59 (2022).

15. Joris, P. X., Schreiner, C. E. & Rees, A. Neural processing of amplitude-modulated sounds. Physiological Reviews 84, 541–577 (2004).

16. Herdman, A. T. et al. Intracerebral sources of human auditory steady-state responses. Brain Topography 15, 69–86 (2002).

17. Kuwada, S. et al. Sources of the scalp-recorded amplitude-modulation following response. J Am Acad Audiol 13, 188–204 (2002).

18. Picton, T. W., John, M. S., Dimitrijevic, A. & Purcell, D. Human auditory steady-state responses. International Journal of Audiology 42, 177–219 (2003).

19. Parthasarathy, A. & Bartlett, E. Two-channel recording of auditory-evoked potentials to detect age-related deficits in temporal processing. Hearing Research 289, p(2012).

20. Shaheen, L. A., Valero, M. D. & Liberman, M. C. Towards a Diagnosis of Cochlear Neuropathy with Envelope Following Responses. J Assoc Res Otolaryngol (2015) doi:10.1007/s10162-015-0539-3.

21. Pichora-Fuller, M. K. et al. Hearing Impairment and Cognitive Energy: The Framework for Understanding Effortful Listening (FUEL). Ear and Hearing 37, 5S–27S (2016).

22. Kahneman, D. & Beatty, J. Pupil diameter and load on memory. Science 154, 1583–(1966).

23. Peelle, J. E. Listening Effort: How the Cognitive Consequences of Acoustic Challenge Are Reflected in Brain and Behavior. Ear and Hearing 39, 204–214 (2018).

24. Beatty, J. Phasic not tonic pupillary responses vary with auditory vigilance performance. Psychophysiology 19, 167–172 (1982).

25. Tun, P. A., McCoy, S. & Wingfield, A. Aging, Hearing Acuity, and the Attentional Costs of Effortful Listening. Psychology and Aging 24, 761–766 (2009).

26. Piquado, T., Isaacowitz, D. & Wingfield, A. Pupillometry as a measure of cognitive effort in younger and older adults. Psychophysiology 47, 560–569 (2010).

27. McGinley, M. J., David, S. V. & McCormick, D. A. Cortical Membrane Potential Signature of Optimal States for Sensory Signal Detection. Neuron 87, 179–192 (2015).

28. Winn, M. B., Edwards, J. R. & Litovsky, R. Y. The Impact of Auditory Spectral Resolution on Listening Effort Revealed by Pupil Dilation. Ear and Hearing 36, e153–e165 (2015).

29. Reimer, J. et al. Pupil fluctuations track rapid changes in adrenergic and cholinergic activity in cortex. Nat Commun 7, 13289 (2016).

30. Killion, M. C., Niquette, P. A., Gudmundsen, G. I., Revit, L. J. & Banerjee, S. Development of a quick speech-in-noise test for measuring signal-to-noise ratio loss in normal-hearing and hearing-impaired listeners. Journal of the Acoustical Society of America 116, 2395–2405 (2004).

31. Nasreddine, Z. S. et al. The Montreal Cognitive Assessment, MoCA: a brief screening tool for mild cognitive impairment. J Am Geriatr Soc 53, 695–699 (2005).

32. Beck, A. T., Steer, R. A. & Carbin, M. G. Psychometric properties of the Beck Depression Inventory: Twenty-five years of evaluation. Clinical Psychology Review 8, 77–100 (1988).

33. Wilson, P. H., Henry, J., Bowen, M. & Haralambous, G. Tinnitus reaction questionnaire: psychometric properties of a measure of distress associated with tinnitus. J Speech Hear Res 34, 197–201 (1991).

34. McHaney, J. R., Tessmer, R., Roark, C. L. & Chandrasekaran, B. Working memory relates to individual differences in speech category learning: Insights from computational modeling and pupillometry. Brain Lang 222, 105010 (2021).

35. McHaney, J. R., Schuerman, W. L., Leonard, M. K. & Chandrasekaran, B. Low amplitude transcutaneous auricular vagus nerve stimulation modulates performance but not pupil size during non-native speech category learning. 2022.07.19.500625 Preprint at https://doi.org/10.1101/2022.07.19.500625 (2022).

36. R Core Team. R: a language and environment for statistical computing. https://www.gbif.org/tool/81287/r-a-language-and-environment-for-statistical-computing (2022).

37. Bates, D., Mächler, M., Bolker, B. & Walker, S. Fitting Linear Mixed-Effects Models Using lme4. Journal of Statistical Software 67, 1–48 (2015).

38. Kuznetsova, A., Brockhoff, P. B. & Christensen, R. H. B. lmerTest Package: Tests in Linear Mixed Effects Models. Journal of Statistical Software 82, 1–26 (2017).

39. Benjamini, Y. & Hochberg, Y. Controlling the False Discovery Rate: A Practical and Powerful Approach to Multiple Testing. Journal of the Royal Statistical Society: Series B (Methodological) 57, 289–300 (1995).

40. David, F. N. Tables of the Ordinates and Probability Integral of the Distribution of the Correlation Coefficient in Small Samples. (Biometrika office, University college, and printed at the University Press, Cambridge, Eng., 1938).

41. Venables, W. N. & Ripley, B. D. Modern Applied Statistics with S. (Springer, 2002). doi:10.1007/978-0-387-21706-2.

42. Galbraith, G. et al. Murine auditory brainstem evoked response: Putative two-channel differentiation of peripheral and central neural pathways. Journal of Neuroscience Methods 153, 214–220 (2006).

43. Ping, J. L. et al. Auditory evoked responses in the rat: Transverse mastoid needle electrodes register before cochlear nucleus and do not reflect later inferior colliculus activity. Journal of Neuroscience Methods 161, 11–16 (2007).

44. Wang, L., Bharadwaj, H. & Shinn-Cunningham, B. Assessing Cochlear-Place Specific Temporal Coding Using Multi-Band Complex Tones to Measure Envelope-Following Responses. Neuroscience 407, 67–74 (2019).

45. Encina-Llamas, G., Dau, T. & Epp, B. On the use of envelope following responses to estimate peripheral level compression in the auditory system. Sci Rep 11, 6962 (2021).

46. Watkins, K. E., Strafella, A. P. & Paus, T. Seeing and hearing speech excites the motor system involved in speech production. Neuropsychologia 41, 989–994 (2003).

47. Shinn-Cunningham, B. G. & Best, V. Selective attention in normal and impaired hearing. Trends Amplif 12, 283–299 (2008).

48. Rönnberg, J., Rudner, M., Lunner, T. & Zekveld, A. A. When cognition kicks in: working memory and speech understanding in noise. Noise Health 12, 263–269 (2010).

49. Golestani, N., Hervais-Adelman, A., Obleser, J. & Scott, S. K. Semantic versus perceptual interactions in neural processing of speech-in-noise. Neuroimage 79, 52–61 (2013).

50. Zekveld, A. A., Rudner, M., Johnsrude, I. S. & Rönnberg, J. The effects of working memory capacity and semantic cues on the intelligibility of speech in noise. J Acoust Soc Am 134, 2225–2234 (2013).

51. Du, Y., Buchsbaum, B. R., Grady, C. L. & Alain, C. Noise differentially impacts phoneme representations in the auditory and speech motor systems. Proc Natl Acad Sci U S A 111, 7126–7131 (2014).

52. Du, Y., Buchsbaum, B. R., Grady, C. L. & Alain, C. Increased activity in frontal motor cortex compensates impaired speech perception in older adults. Nat Commun 7, 12241 (2016).

53. McGarrigle, R., Dawes, P., Stewart, A. J., Kuchinsky, S. E. & Munro, K. J. Pupillometry reveals changes in physiological arousal during a sustained listening task. Psychophysiology 54, 193–203 (2017).

54. Kousaie, S. et al. Language learning experience and mastering the challenges of perceiving speech in noise. Brain Lang 196, 104645 (2019).

55. Price, C. N. & Bidelman, G. M. Attention reinforces human corticofugal system to aid speech perception in noise. NeuroImage 235, 118014 (2021).

56. Holmes, E., Purcell, D. W., Carlyon, R. P., Gockel, H. E. & Johnsrude, I. S. Attentional Modulation of Envelope-Following Responses at Lower (93–109 Hz) but Not Higher (217– 233 Hz) Modulation Rates. JARO 19, 83–97 (2018).

57. Shinn-Cunningham, B. Cortical and Sensory Causes of Individual Differences in Selective Attention Ability Among Listeners With Normal Hearing Thresholds. J Speech Lang Hear Res 60, 2976–2988 (2017).

58. DiNino, M., Holt, L. L. & Shinn-Cunningham, B. G. Cutting Through the Noise: Noise-Induced Cochlear Synaptopathy and Individual Differences in Speech Understanding Among Listeners With Normal Audiograms. Ear Hear 43, 9–22 (2022).

59. Mamo, S. K. & Helfer, K. S. Speech Understanding in Modulated Noise and Speech Maskers as a Function of Cognitive Status in Older Adults. Am J Audiol 30, 642–654 (2021).

60. Xie, Z., Zinszer, B. D., Riggs, M., Beevers, C. G. & Chandrasekaran, B. Impact of depression on speech perception in noise. PLoS One 14, e0220928 (2019).

61. Smayda, K. E., Engen, K. J. V., Maddox, W. T. & Chandrasekaran, B. Audio-Visual and Meaningful Semantic Context Enhancements in Older and Younger Adults. PLOS ONE 11, e0152773 (2016).

62. Grant, K. J. et al. Predicting neural deficits in sensorineural hearing loss from word recognition scores. Sci Rep 12, 8929 (2022).

63. Holmes, E. & Griffiths, T. D. ‘Normal’ hearing thresholds and fundamental auditory grouping processes predict difficulties with speech-in-noise perception. Sci Rep 9, 16771 (2019).

64. Picton, T. W., Skinner, C. R., Champagne, S. C., Kellett, A. J. C. & Maiste, A. C. Potentials-evoked by the sinusoidal modulation of the amplitude or frequency of a tone. Journal of the Acoustical Society of America 82, 165–178 (1987).

65. Boettcher, F. A., Poth, E. A., Mills, J. H. & Dubno, J. R. The amplitude-modulation following response in young and aged human subjects. Hearing Research 153, 32–42 (2001).

66. He, N. J., Mills, J. H., Ahlstrom, J. B. & Dubno, J. R. Age-related differences in the temporal modulation transfer function with pure-tone carriers. Journal of the Acoustical Society of America 124, 3841–3849 (2008).

67. Ruggles, D., Bharadwaj, H. & Shinn-Cunningham, B. G. Normal hearing is not enough to guarantee robust encoding of suprathreshold features important in everyday communication. Proceedings of the National Academy of Sciences of the United States of America 108, 15516–15521 (2011).

68. Bharadwaj, H. M., Masud, S., Mehraei, G., Verhulst, S. & Shinn-Cunningham, B. G. Individual Differences Reveal Correlates of Hidden Hearing Deficits. Journal of Neuroscience 35, 2161–2172 (2015).

69. Dimitrijevic, A. et al. Human Envelope Following Responses to Amplitude Modulation: Effects of Aging and Modulation Depth. Ear and Hearing 37, E322–E335 (2016).

70. Chandrasekaran, B. & Kraus, N. The scalp-recorded brainstem response to speech: Neural origins and plasticity. Psychophysiology 47, 236–246 (2010).

71. Coffey, E. B. J., Herholz, S. C., Chepesiuk, A. M. P., Baillet, S. & Zatorre, R. J. Cortical contributions to the auditory frequency-following response revealed by MEG. Nat Commun 7, 11070 (2016).

72. Coffey, E. B. J. et al. Evolving perspectives on the sources of the frequency-following response. Nat Commun 10, 5036 (2019).

73. Bidelman, G. M. Subcortical sources dominate the neuroelectric auditory frequency-following response to speech. Neuroimage 175, 56–69 (2018).

74. Gnanateja, G. N. et al. Frequency-Following Responses to Speech Sounds Are Highly Conserved across Species and Contain Cortical Contributions. eNeuro 8, ENEURO.0451-21.2021 (2021).

75. Kiren, T., Aoyagi, M., Furuse, H. & Koike, Y. An experimental-study on the generator of amplitude-modulation following response. Acta Oto-Laryngologica 28–33 (1994).

76. Mepani, A. M. et al. Envelope following responses predict speech-in-noise performance in normal-hearing listeners. J Neurophysiol 125, 1213–1222 (2021).

77. Galbraith, G. C. 2-Channel brain-stem frequency-following responses to pure-tone and missing fundamental stimuli. Electroencephalography and Clinical Neurophysiology 92, 321–330 (1994).

78. Galbraith, G. C. et al. Putative measure of peripheral and brainstem frequency-following in humans. Neuroscience Letters 292, 123–127 (2000).

79. King, A., Hopkins, K. & Plack, C. J. Differential Group Delay of the Frequency Following Response Measured Vertically and Horizontally. Jaro-Journal of the Association for Research in Otolaryngology 17, 133–143 (2016).

80. Wild, C. J. et al. Effortful Listening: The Processing of Degraded Speech Depends Critically on Attention. Journal of Neuroscience 32, 14010–14021 (2012).

81. Kuchinsky, S. E. et al. Pupil size varies with word listening and response selection difficulty in older adults with hearing loss. Psychophysiology 50, 23–34 (2013).

82. Kuchinsky, S. E. et al. Task-Related Vigilance During Word Recognition in Noise for Older Adults with Hearing Loss. Exp Aging Res 42, 50–66 (2016).

83. Winn, M. B. Rapid Release From Listening Effort Resulting From Semantic Context, and Effects of Spectral Degradation and Cochlear Implants. Trends in Hearing 20, p(2016).

84. McLaughlin, D. J. et al. Give me a break! Unavoidable fatigue effects in cognitive pupillometry. Psychophysiology e14256 (2023) doi:10.1111/psyp.14256.

85. Aston-Jones, G. & Cohen, J. D. An integrative theory of locus coeruleus-norepinephrine function: Adaptive gain and optimal performance. Annual Review of Neuroscience 28, 403–450 (2005).

86. de Gee, J. W. et al. Pupil-linked phasic arousal predicts a reduction of choice bias across species and decision domains. Elife 9, e54014 (2020).

87. McGinley, M. J. et al. Waking State: Rapid Variations Modulate Neural and Behavioral Responses. Neuron 87, 1143–1161 (2015).

